# Asymmetric sampling in human auditory cortex reveals spectral processing hierarchy

**DOI:** 10.1101/581520

**Authors:** Jérémy Giroud, Agnès Trébuchon, Daniele Schön, Patrick Marquis, Catherine Liegeois-Chauvel, David Poeppel, Benjamin Morillon

**Affiliations:** Aix Marseille Univ, Inserm, INS, Inst Neurosci Syst, Marseille, France; Cleveland Clinic Neurological Institute, Epilepsy Center, Cleveland, OH, USA; Department of Neuroscience, Max-Planck-Institute for Empirical Aesthetics, Frankfurt am Main, Germany; Department of Psychology and Center for Neural Science, New York University, New York, NY, USA

**Keywords:** hemispheric asymmetry, intracranial EEG, speech, neural oscillations, functional cytoarchitectonic

## Abstract

Speech perception is mediated by both left and right auditory cortices, but with differential sensitivity to specific acoustic information contained in the speech signal. A detailed description of this functional asymmetry is missing, and the underlying models are widely debated. We analyzed cortical responses from 96 epilepsy patients with electrode implantation in left or right primary, secondary, and/or association auditory cortex. We presented short acoustic transients to reveal the stereotyped spectro-spatial oscillatory response profile of the auditory cortical hierarchy. We show remarkably similar bimodal spectral response profiles in left and right primary and secondary regions, with preferred processing modes in the theta (∼4-8 Hz) and low gamma (∼25-50 Hz) ranges. These results highlight that the human auditory system employs a two-timescale processing mode. Beyond these first cortical levels of auditory processing, a hemispheric asymmetry emerged, with delta and beta band (∼3/15 Hz) responsivity prevailing in the right hemisphere and theta and gamma band (∼6/40 Hz) activity in the left. These intracranial data provide a more fine-grained and nuanced characterization of cortical auditory processing in the two hemispheres, shedding light on the neural dynamics that potentially shape auditory and speech processing at different levels of the cortical hierarchy.

**Author summary:** Speech processing is now known to be distributed across the two hemispheres, but the origin and function of lateralization continues to be vigorously debated. The asymmetric sampling in time (AST) hypothesis predicts that (1) the auditory system employs a two-timescales processing mode, (2) present in both hemispheres but with a different ratio of fast and slow timescales, (3) that emerges outside of primary cortical regions. Capitalizing on intracranial data from 96 epileptic patients we sensitively validated each of these predictions and provide a precise estimate of the processing timescales. In particular, we reveal that asymmetric sampling in associative areas is subtended by distinct two-timescales processing modes. Overall, our results shed light on the neurofunctional architecture of cortical auditory processing.

## Introduction

Contrary to the classic neuropsychological perspective, speech processing is now known to be distributed across the two hemispheres, with some models positing a leftward dominance for verbal comprehension and a rightward dominance for processing suprasegmental features, including aspects of prosody or voice processing [1]. The origin and function of lateralization continues to be vigorously debated, for example with regard to its domain-general or domain-specific nature [2,3]. The former view predicts that lateralization of speech processing (and auditory processing, in general) originates in general-purpose mechanisms sensitive to the low-level acoustic features present in speech. The domain-specific view postulates that speech is processed in a dedicated system lateralized to the left hemisphere. On this view, processing critically depends on the specific linguistic properties of a stimulus. Crucial to this debate is, thus, proper understanding of the distinctive sensitivity of the left and right auditory cortical regions to acoustic features, which should be grounded in characteristic anatomic-functional signatures.

There exists suggestive neuroanatomical evidence for structural differences between the left and right auditory cortex. The density of white and grey matter is higher in the left hemisphere, respectively in Heschl’s gyrus and in both the planum temporale and Heschl’s gyrus [4,5]. Moreover, the left auditory cortex contains larger cortical columns with a higher number of large pyramidal cells in cortical layer III than its right counterpart [6]. Those differences in cytoarchitectonic organization should coexist with electrophysiological and functional differences between auditory regions. Building on such observations, the asymmetric sampling in time (AST) hypothesis makes several interrelated predictions related to the characteristics of auditory information processing at the cortical level [7]:

(1) The human auditory system employs (at least) a two-timescale processing mode, characterized by oscillatory cycles that can be viewed as individual computational units. These two timescales operate in the low-gamma (∼25-50 Hz) and theta (∼4-8 Hz) frequency ranges, corresponding, respectively, to a temporal integration window of ∼30 ms and ∼200 ms. Such temporal multiplexing allows the system to process in parallel acoustic information using two complementary algorithmic strategies, optimized to encode complementary spectrotemporal characteristic of sounds. This prediction - that sounds are processed at preferred and specific timescales - has received support from both auditory and speech-specific paradigms [8–16].

(2) This dual-timescale processing operates in both hemispheres, but the ratio of neural ensembles dedicated to the processing of each timescale differs between left and right hemispheres. Indeed, while the left auditory cortex would preferentially process auditory streams using a short temporal integration window (∼30 ms), the right auditory cortex would preferentially sample information using a long temporal integration window (∼200 ms). Previous findings reported that left and right cortical auditory regions exhibit differences in their intrinsic oscillatory activity [17–19]. A relative leftward dominance of low-gamma neural oscillations and/or rightward dominance of theta oscillations is also visible during sensory stimulation [18,20,21]. This asymmetry is, moreover, reflected in the sensitivity of the left and right auditory cortex to different spectrotemporal modulations of sounds, with a leftward dominance for fast temporal modulations, and/or a rightward dominance for slow temporal modulations [13,22–29].

(3) The electrophysiological signature of this asymmetry emerges outside of primary auditory regions. The AST hypothesis in its original conception posited that at the level of core auditory cortex there is no obvious functional asymmetry, but that beyond this first stage of cortical processing a functional asymmetry should be visible, namely in left and right association auditory regions. This last point has also received some empirical support [13,25,26,28].

While each of these predictions has received experimental support, they are also vigorously debated. In particular, one concern relates to the specificity of the left temporal lobe for faster temporal modulations. Some authors have suggested that most published results can be interpreted in an alternative framework, wherein only the right temporal lobe shows a marked preference for certain properties of sounds, for example longer durations, or variations in pitch [3,30]. Moreover, by contrast with the AST hypothesis, some authors suggested that the hemispheric asymmetry may stem from core auditory areas (Heschl gyrus) and not association cortex [17–19,22,23,31–33]. The conflicting results may be due to differences in paradigms, stimuli, as well as resolution of the imaging instruments employed. Discrepancies may also arise from the fact that this asymmetry probably takes different forms along the auditory cortical pathway, with a more subtle functional signature at early cortical stages and more striking qualitative differences at later processing stages (see e.g., [18]). Finally, and this is a crucial aspect of the theory, the duration of these temporal integration windows was never precisely characterized physiologically with high-resolution data.

To adjudicate between the models, more sensitively test the predictions, and overcome some of the difficulties in acquiring decisive data to characterize the signature of auditory hemispheric lateralization at both high spatial and temporal (hence spectral) resolutions, we combined two novel experimental approaches. (1) We capitalize on a recent methodology developed to investigate the functional cytoarchitectonics of auditory regions, which consists of presenting transient acoustic impulses to probe a stereotyped response function at the neural level, revealing the intrinsic network dynamic properties of the recorded neural population [34]. This method allows us to measure the preferred oscillatory modes of recorded neural ensembles. (2) We capitalize on data acquired from 96 epileptic patients, implanted for clinical evaluation at various stages of the auditory cortical hierarchy. Our results show the natural spectral profile of neural activity in left and right primary, secondary, and association (anterior BA22) cortical auditory regions, thus enabling a detailed characterization of the *potential* inter-hemispheric functional differences and dynamics at play during auditory processing.

## Results

Data from 96 epileptic patients implanted with depth macroelectrodes located in left and right primary (PAC) secondary (SAC) and association (AAC) auditory cortex were analysed (Fig. 1) [35–38]. PAC, SAC, and AAC respectively correspond to the postero-medial portion of Heschl’s gyrus (A1, medial belt and lateral belt areas), the lateral posterior superior temporal gyrus (STG; parabelt area) and the lateral anterior STG (area A4) [39]. Patients participated in a perceptual experiment during which they passively listened to pure tones and syllables (see Materials and Methods).

**Figure 1:**
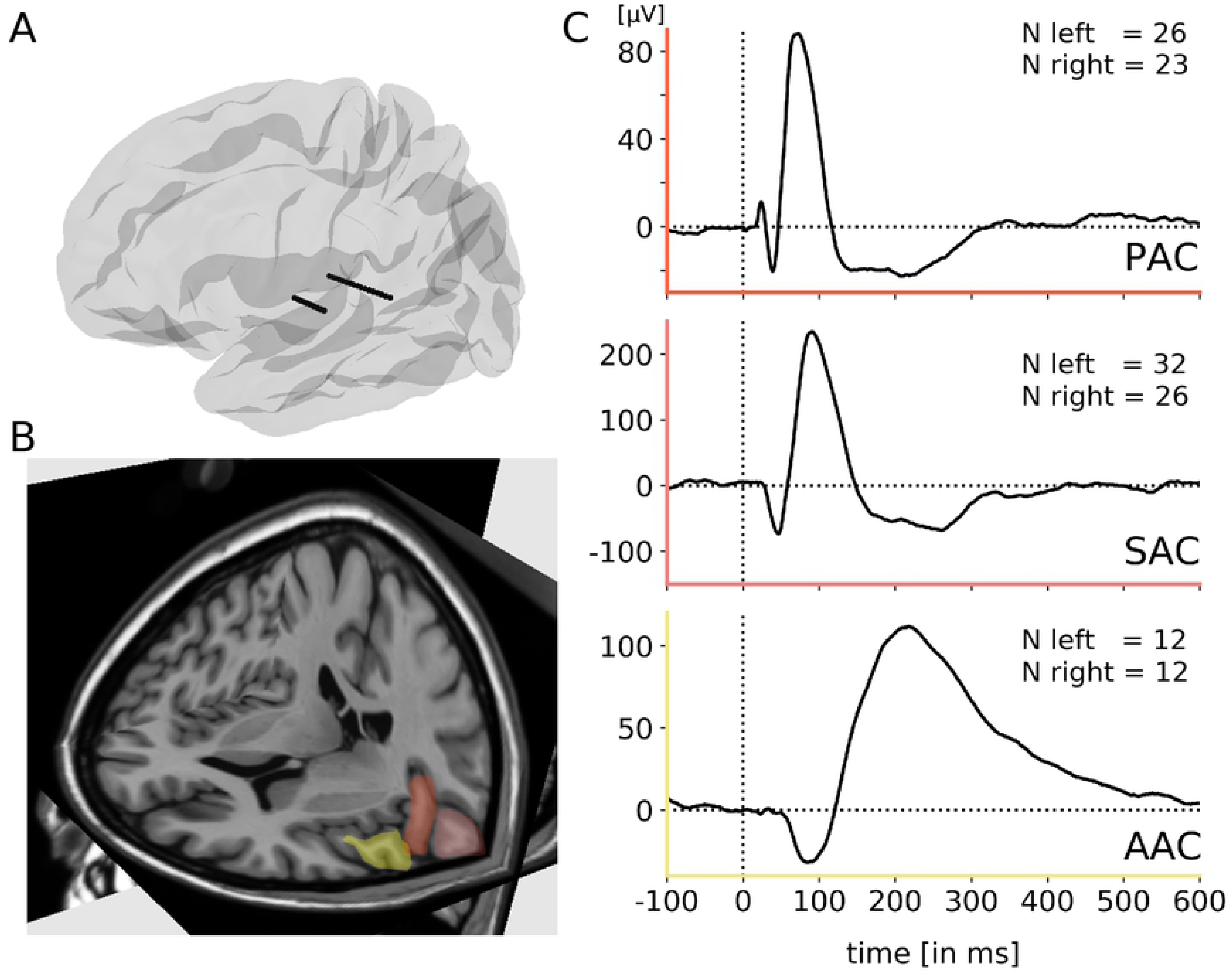
Example of electrodes and contact positions with characteristic auditory evoked potentials (AEPs) in a patient implanted in the left hemisphere. **(A)** Example of typical SEEG recordings electrodes shown in a 3D view of the left hemisphere, on a template brain in Montreal Neurological Institute space. The posterior electrode is composed of 15 contacts and targets core auditory regions, while the anterior electrode, composed of 10 contacts, targets association auditory regions. **(B)** MRI scan showing the location of the 3 regions of interest (ROIs) in the left hemisphere: i) Primary auditory cortex (PAC): the postero-medial portion of Heschl’s gyrus (A1, medial belt and lateral belt areas; see [39]) in red; ii) Secondary auditory cortex (SAC): the lateral posterior superior temporal gyrus (STG; parabelt area) in pink; iii) Association auditory cortex (AAC): the lateral anterior STG (area A4) in yellow. **(C)** AEPs in response to pure tones from a representative patient, for each ROI. The axes are color-coded according to the locations displayed in B. Electrode contacts used along the shaft were selected based on their anatomical location and functional responses (typical shape and latencies of evoked responses; see Materials and Methods). The insets indicate the number of patients recorded at each location.

### Spectral characteristics of the evoked response to transient pure tones

To investigate the fine-grained temporal constraints of the first cortical stages of the auditory processing hierarchy, we first analysed the evoked responses to transient acoustic impulses (30 ms duration pure tones, presented at 0.5 or 1 kHz). A time-frequency representation of the evoked responses as computed through inter-trial coherence (ITC) (Fig. 2A) demonstrates the presence of multiple co-occurring stereotyped (i.e. identical across trials) spectral components, either within or between regions of interest (ROIs). These components were limited in time and homogenous, and their central frequency was best captured by averaging ITC values over time (see Materials and Methods). In PAC and SAC, a group-level analysis revealed the presence of a single stereotyped response profile, characterized by a main spectral maximum within the theta range (∼4-8 Hz; corresponding to a time-constant of ∼150 ms; Fig. 2B). Importantly, this response profile was similar across left and right hemispheres. Conversely, a more complex pattern of response was visible in AAC, with the presence of two distinct salient spectral maxima that differed between left and right hemispheres. Prominent spectral peaks in the theta (4-8 Hz) and gamma (25-45 Hz) frequency ranges were visible in left AAC; the right counterpart was characterized by peaks in the delta (1-4 Hz) and beta (15-30 Hz) frequency ranges (Fig. 2B).

**Figure 2:**
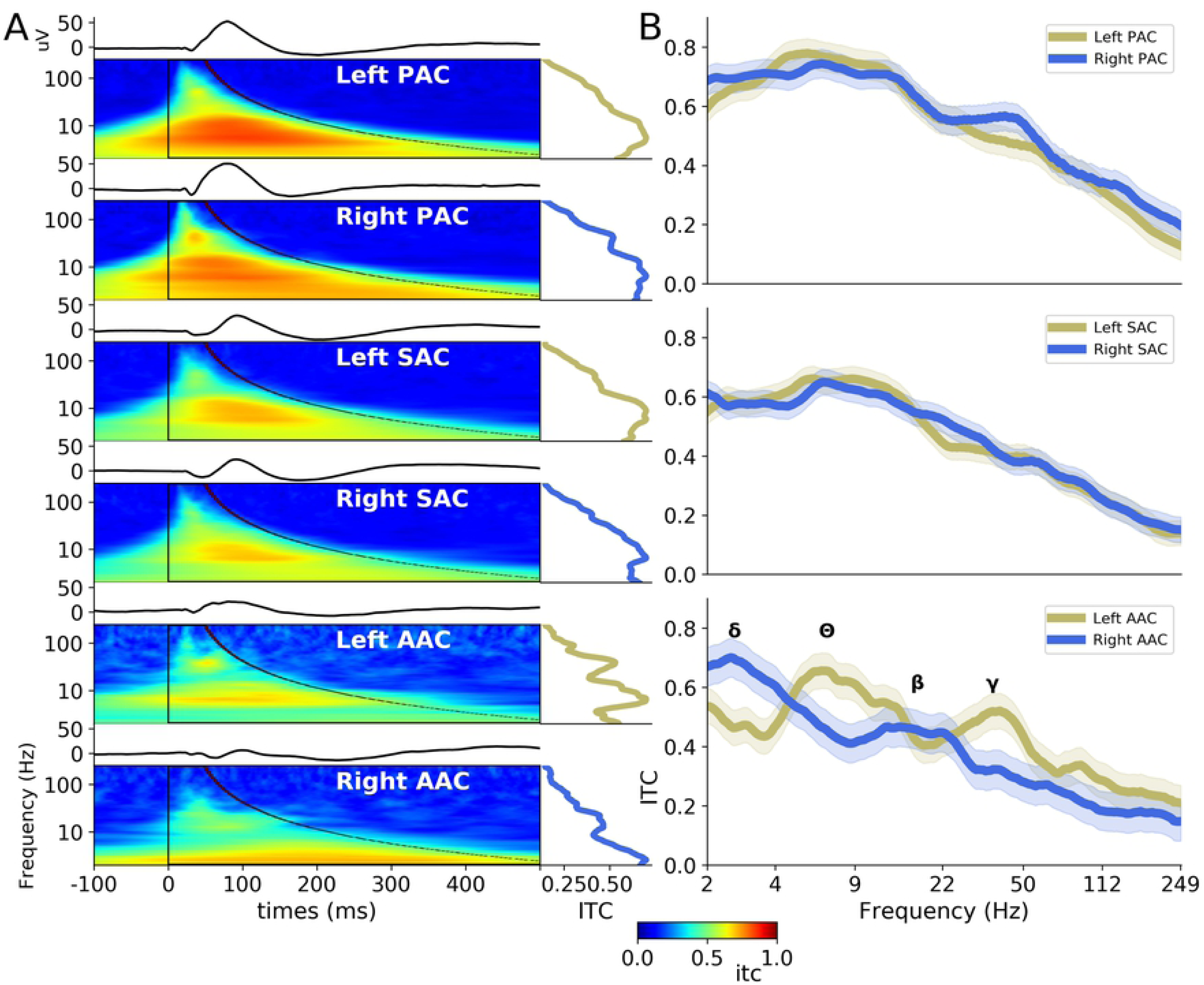
Evoked activity in response to pure tones (0.5 kHz and 1 kHz tone responses merged) in hierarchically organized auditory areas. **(A)** AEPs (top panels) and intertrial coherence (ITC; lower panels) in response to pure tones, averaged across participants, in the six auditory regions of interest (PAC, SAC and AAC, in left and right hemispheres). Right insets indicate the spectrum of the ITC, averaged over the window of interest (as the time varies with the frequency) (black overlay) **(B)** Inter-hemispheric comparison of the ITC spectra in response to pure tones, for PAC, SAC and AAC. Shaded areas indicate SEM. Greek letters indicate the main peaks observed in AAC ([Symbol]: delta 1-4 Hz; [Symbol]: theta 4-8 Hz; [Symbol]: beta 14-30 Hz; [Symbol]: low-gamma 25-45 Hz).

To better characterize the time-constants of the neural processes occurring at each putative step of the auditory cortical hierarchy, we extracted for each participant and ROI the two highest local maxima of the ITC spectrum (between 2-250 Hz; Fig. 3). This analysis substantiates the finding that in PAC and SAC the evoked response was dominated by a spectral peak in the theta range (∼4-8 Hz), and highlights that a secondary peak emerged in the gamma range (∼25-50 Hz). At these earlier cortical stages, no differences in the spectral profile of the transient evoked activity emerged across hemispheres, neither in PAC (Mann Whitney U test: 1st peak, U = 222.0, p = .09; 2nd peak, U = 251.5, p = .24), nor SAC (1st peak, U = 380.5, p = .29; 2nd peak, U = 373.5, p = .25). In contrast, in AAC a more complex and significantly asymmetric spectral profile emerged. The evoked response was characterized by faster components in left than right AAC, with respectively a theta/gamma (∼6/40 Hz) spectral profile in left and a delta/beta (∼3/15 Hz) spectral profile in right AAC. Inter-hemispheric comparison of the spectral distribution of the two main ITC peaks confirmed that this asymmetry was significant (1st peak, U = 0.0, *p* < 0.001; 2nd peak, U = 0.0, *p* < .001).

**Figure 3.**
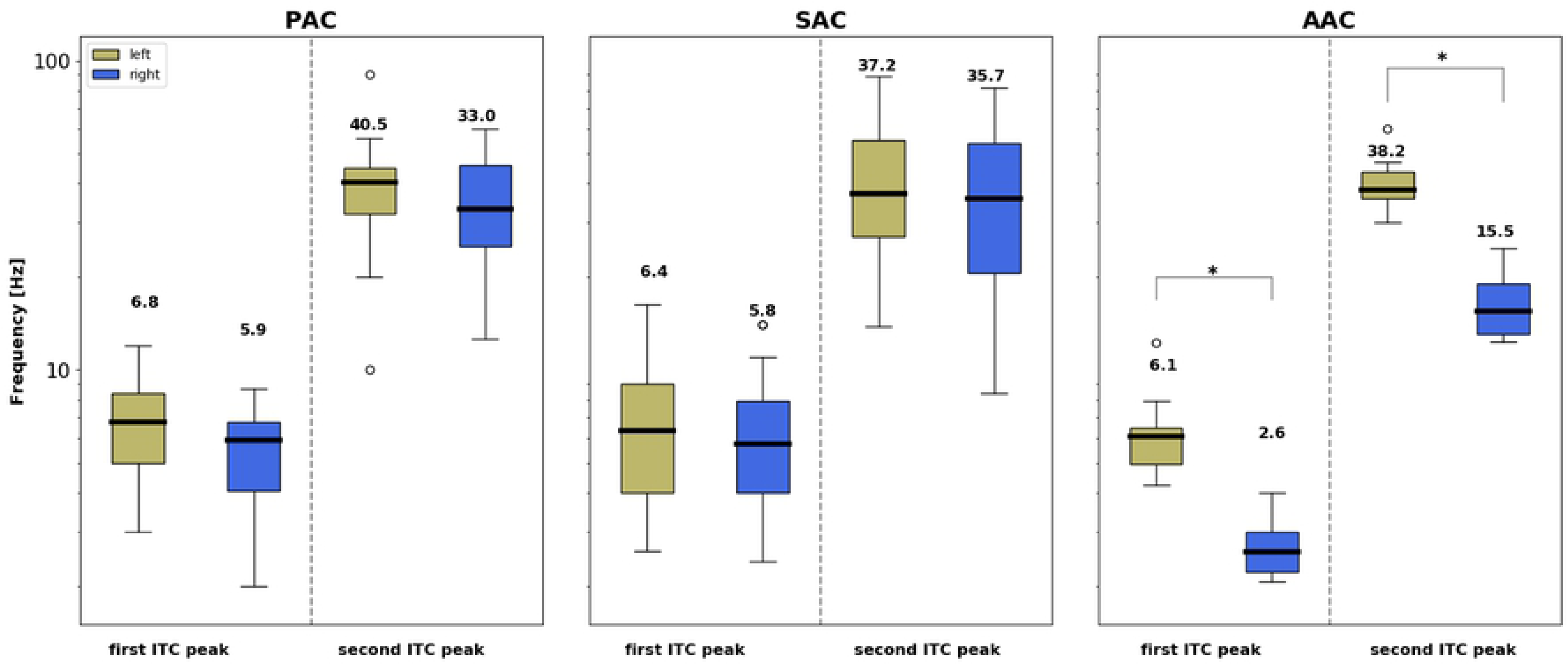
Inter-hemispheric comparison of the spectral distribution of the two main ITC peaks in response to pure tones (0.5 kHz and 1 kHz tone responses merged) in the different auditory areas. Individual peaks were identified as the two highest local maxima of the ITC spectrum (between 2-250 Hz). Frequency (in Hz) is denoted on the y-axis. Box plots: The centre line within each box denotes the median frequency, and the bottom and top of each box designate the first and third quartile, respectively. Number of patients recorded at each location: PAC left = 26, PAC right = 23, SAC left = 32, SAC right = 26, AAC left = 12, and AAC right = 12. * indicates significant inter-hemispheric differences (Mann Whitney U Test, p < .01).

Next, to confirm the robustness of these findings, we reanalysed the evoked response to pure tones separately for 0.5 and 1 kHz pure tones, and observed the exact same ITC spectral profile for each ROI, independently of the frequency of the pure tone (Fig. S1). The statistical analyses also confirmed the absence of significant inter-hemispheric differences in PAC and SAC (all U > 280.0, all *p* > 0.08), and the emergence of an asymmetry in AAC (0.5 kHz: 1st peak, U = 1.0, p < 0.001; 2nd peak, U = 4.0, p < .001; 1 kHz: 1st peak, U = 0.5, p < 0.001; 2nd peak, U = 0.5, p <0 .001).

### Interaction between stimulus and neural dynamics

Finally, the same analysis was carried on data recorded on the same patients during presentation of syllables (French /ba/ and /pa/; Fig. 4). Importantly, these stimuli are characterized by more complex spectro-temporal dynamics than transient pure tones. Accordingly, we observed that the spectral profile of the ITC differed between pure tones, /ba/ and /pa/ stimuli, as predicted.

**Figure 4:**
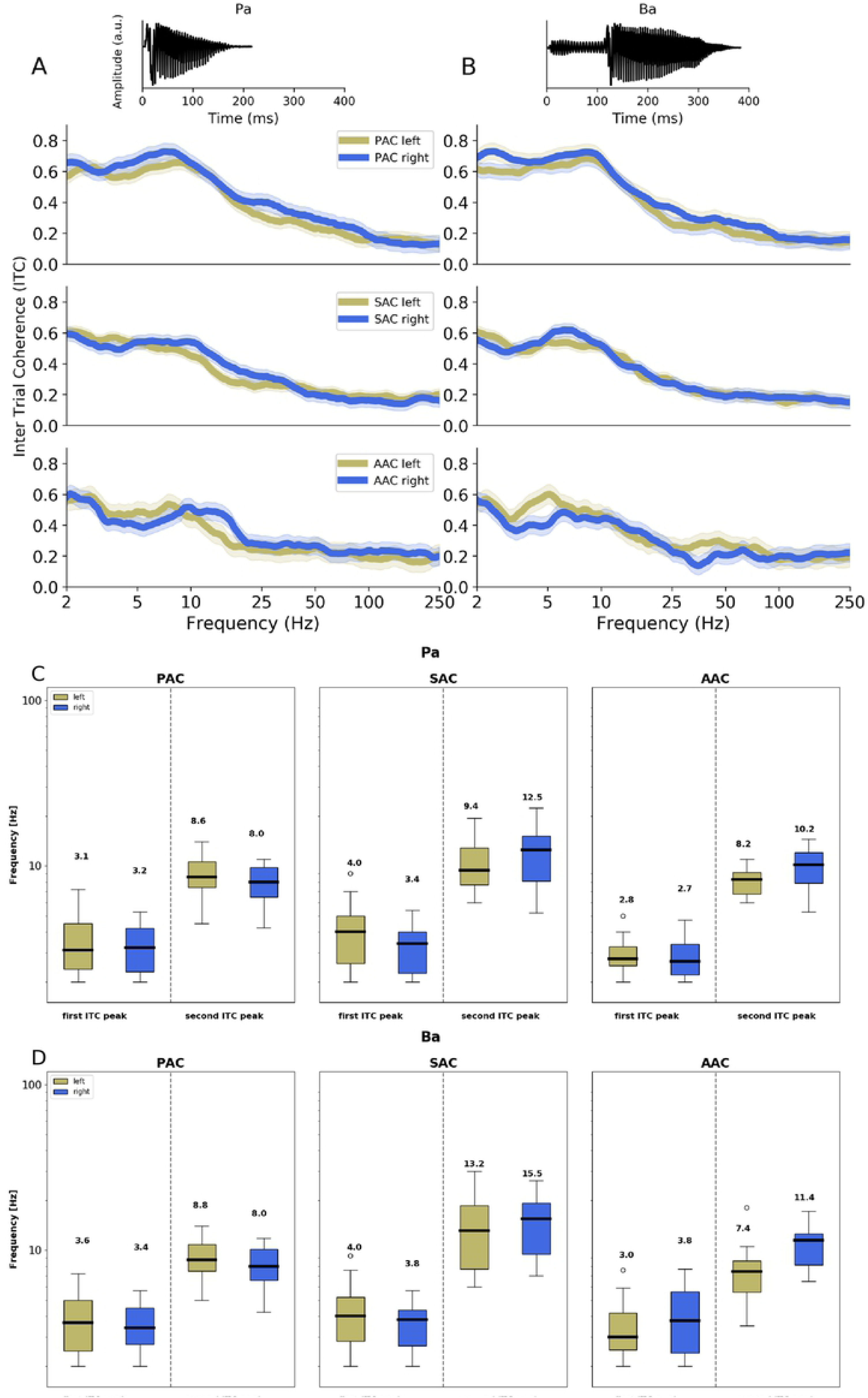
Evoked activity in response to French syllables /Pa/ and /Ba/ in auditory areas. ***(A-B)** Top panels: acoustic waveform of the syllables. Lower panels: Inter-hemispheric comparison of the ITC spectra in response to the syllables* **(A)** /Pa/ and **(B)** /Ba/, *for PAC, SAC and AAC. Shaded areas indicate SEM across participants. **(C-D)*** Inter-hemispheric comparison of the spectral distribution of the two main ITC peaks in response to French syllables **(C)** /Pa/ and **(D)** /Ba/, in auditory areas. Same conventions than in Figure 3.

Results with syllables yielded less prominent and specific spectral peaks in the different ROIs, even in the latter stages of auditory processing (AAC). The maximum neural activity in response to syllable presentation was in the low frequency range (<20Hz) and did not change across regions of interest.

## Discussion

Derived from intracranial recordings from 96 epileptic patients, and using transient auditory stimulation with tones and syllables, this study aimed to characterize the intrinsic timescales of auditory information processing at three stages of the auditory cortical hierarchy. The spatial and temporal precision offered by SEEG recording enables a meticulous description of the neural dynamics at each anatomic stage. Using transient acoustic stimulation allowed us to probe stereotyped neural responses which reveal the intrinsic organizational dynamics of the recorded areas.

Our results reveal, first, that the early cortical stages of auditory processing (PAC, SAC), when acoustically stimulated with transient stimuli, show characteristic bimodal spectral profiles. These functional responses were characterized by components in the theta (∼4-8 Hz) and gamma (∼25-50 Hz) ranges. This result, obtained with high-resolution data on a large cohort of patients is consistent with previous findings obtained with intracranial recordings on a single case [19]. Moreover, the presence of two concomitant oscillatory modes is a strong evidence in favor of the AST framework, in which the auditory system makes use of a two-timescale processing mode to perceptually sample acoustic dynamics [10,14]. This specific bimodal neural signature also corroborates physiological descriptions of a natural interaction between low and high oscillatory frequencies (phase-amplitude coupling) at rest and during auditory stimulation at the level of local neural ensembles [40]. No hemispheric difference is apparent in core auditory regions, as evidenced by the similar bimodal spectral pattern elicited in left and right PAC and SAC. This is in contradistinction with previous findings describing functional asymmetries in core auditory areas [17–19,22,23,31–33]. One limitation of SEEG recording is the absence of whole brain coverage. In particular, our functional characterization of early cortical auditory processing was limited to the postero-medial portion of Heschl’s gyrus (PAC) and the lateral posterior STG (SAC; Fig. 1). We hence cannot exclude the presence of a functional asymmetry in core auditory regions, notably in the lateral portion of Heschl’s gyrus and the planum polare [41], which we did not sample. However, our results are compatible with current models of the functional organisation of the core auditory cortex, which report relatively weak functional hemispheric differences [42]. Second, our results show the emergence of a strong functional asymmetry at the level of the association auditory cortex (AAC). We observed prominent spectral peaks in theta and gamma bands (∼6/40 Hz) in the left AAC, whereas its right counterpart was characterized by delta and beta (∼3/15 Hz) oscillatory modes. Of note, all the areas investigated were characterized by a bimodal spectral profile. However, while most of these results are in accordance with the AST – i.e. observation of a bimodal spectral profile; emergence of functional asymmetry in association areas – they also reveal that this functional asymmetry does not simply correspond to a differential ratio of neural ensembles oscillating at theta and gamma rates [7], but in fact correspond to the involvement of distinct oscillatory modes (theta/gamma vs. delta/beta) between left and right hemispheres.

The spectral profile of neural response in the left AAC (but also bilateral PAC and SAC) can be linked to a recent model of coupled oscillators describing the sensory analysis of speech, in which low and high frequency oscillations operate in the theta and gamma ranges, respectively, and process in parallel acoustic information [43]. In this model, the tracking of slow speech fluctuations by theta oscillations and its coupling to gamma activity both appear as critical features for accurate speech encoding, underscoring the importance of a two-timescale processing mode for efficiently analysing speech. Moreover, this model suggests that during speech perception syllabic-and phonemic-scale computations operate in combination at a local cortical level of processing, which could correspond to the left AAC.

On the other hand, the presence of neural activity in the delta and beta ranges in right AAC is more puzzling. Previous studies claimed that parsing at the syllabic scale occurs bilaterally [44], or is even rightward lateralized [15,24]. However, the right auditory cortex is more sensitive to spectral than temporal modulations [28,29,45] and perception of prosody is a right-lateralized process [46]. Thus, our observation of a specific oscillatory mode of processing in right AAC could reflect neural mechanisms dedicated to the parsing of spectral acoustic dynamics. Prosodic phenomena at the level of intonation contours are an example of such a phenomenon; the successful perceptual analysis of spoken language requires the processing of the rhythmic and melodic variations in speech to gain knowledge about speaker’s emotions and intentions [46]. The delta intrinsic time-scale observed in right AAC would be particularly well suited to the segmentation of prosodic cues, as they naturally unfold at 0.5-3 Hz, as also argued in a recent computational model [47].

Overall, our results shed light on the neurofunctional architecture of cortical auditory processing, and in particular on the specific processing timescales of different cortical areas. These general mechanisms are thought to apply to general auditory as well as speech perception. They would suggest that syllabic and phonemic information is segmented in parallel locally, through coupled theta and gamma oscillations, while right-lateralized processes, such as intonation contour or prosody perception, would be segmented by delta (and beta) oscillations. The methodology we employ here is only suited to transient stimuli, as longer stimuli – with a temporal dynamics – impose strong temporal constraints on the neural activity, and the resulting dynamics correspond to a more heterogeneous and elaborate interaction between the acoustic features of the stimulus and the intrinsic neural activity. It is thus an ‘intermediate’ method between resting state and speech paradigms, allowing a more precise description of the natural dynamics at play throughout the auditory pathway.

## Materials and methods

### Participants

96 patients (44 females) with pharmacoresistant epilepsy took part in the study. They were implanted with depth electrodes for clinical purpose at the Hôpital de La Timone (Marseille). Their native language was French. Neuropsychological assessments carried out before stereotactic EEG (SEEG) recordings indicated that all participants had intact language functions and met the criteria for normal hearing. None of them had their epileptogenic zone including the auditory areas as identified by experienced epileptologists. Patients provided informed consent prior to the experimental session, and the study was approved by the Institutional Review board of the French Institute of Health (IRB00003888).

### Stimuli and paradigm

Two types of auditory stimuli were presented to the participants in two separate sessions: 1. 30-ms long pure tones, presented binaurally at 500 Hz or 1 kHz (with a linear rise and fall time of 0.3 ms) 110 times each, with an ISI of 1030 (±200) ms; and 2. /ba/ or /pa/ syllables, pronounced by a French female speaker (Fig. 4, top) and presented binaurally 250 times each, with an ISI of 1030 (±200) ms. During the two recording sessions, participants laid comfortably in a chair in a sound attenuated room and listen passively to the stimuli. Auditory stimuli were delivered from loudspeakers in front of the participants at a comfortable volume. Stimuli were presented in a pseudorandom order at a 44 kHz rate using E-prime 1.1 (Psychology Software Tools Inc., Pittsburgh, PA, USA).

### Anatomo-functional definition of contact positions

Depth electrodes (0.8 mm; Alcis, Besançon, France) containing 10 to 15 contacts were used to perform the functional stereotactic exploration. Contacts were 2 mm long and spaced from each other by 1.5 mm. The locations of the electrode implantations were determined solely on clinical grounds. The anatomical position of the contacts of interest for the study was established based on a scanner image acquired after electrodes’ implantation. For each participant, auditory evoked potentials (AEPs) in response to pure tones (500 Hz and 1 kHz merged) were epoched between -5 s to 5 s relative to stimulus onset. Epochs with artefacts and epileptic spikes were discarded by visual inspection, prior to being averaged over trials. A baseline correction was applied on each trial by computing a z-score relative to activity during the baseline from 500 ms to 50 ms before stimulus onset.

AEPs served as a functional indicator to determine the different auditory areas [35,36]. The regions of interest (ROIs) were functionally defined based on the presence of specific components (early P20/N30, N/P50, and N/P 60-100) for primary auditory cortex (PAC), secondary auditory cortex (SAC) and association auditory cortex (AAC) respectively [48]. Among the 96 participants, respectively 26, 32 and 12 had (at least) a contact in left PAC, SAC and AAC, and 23, 26 and 12 had a contact in right PAC, SAC and AAC (Fig. 1). For each participant and ROI, the most responsive contact (i.e., the contact with the largest AEP) was selected for subsequent analyses when multiple contacts were present in the functional ROI.

### SEEG recordings and analysis

SEEG signals were recorded at a sampling rate of 1000 Hz using a 256-channel BrainAmp amplifier system (Brain Products GmbH, Munich, Germany) and bandpass filtered between 0.3 and 500Hz. A scalp electrode placed in Fz was used as the recording reference. SEEG data were epoched between -5 s to 5 s relative to stimulus onset (either pure tones or syllables). Such a long temporal window for epoching allowed a more precise frequency resolution for time frequency analysis. Epochs with artefacts and epileptic spikes were discarded by visual inspection. Data were referenced into a bipolar montage, by subtracting activity recorded at each contact of interest from activity acquired at its closest neighbour site within the same electrode.

Trial-by-trial time-frequency analysis was carried out in a frequency range of 2 to 250 Hz (logarithmically spaced). Morlet wavelet transform was applied to the data using the MNE-python function *time_frequency.tfr_morlet* (n_cycles = 7 and frequency steps = 100) [49]. The frequency-resolved evoked activity was computed across trials by means of the intertrial coherence (ITC). For each time and frequency point, an ITC value close to 0 reflects low consistency of phase angles across trials, whereas an ITC value of 1 reflects a perfect consistency of phase angles across trials. The ITC spectrum was then computed by averaging ITC values within a time-window of interest designed to encompass one oscillatory cycle per frequency (e.g., 0-500 ms at 2 Hz; 0-20 ms at 50 Hz).

To investigate the frequency at which the evoked activity was maximal and be able to compare them across ROI and participants, the resulting ITC spectra were normalized (z-scored) across frequencies. Finally, the two main peaks (highest non-contiguous local maxima) of the ITC spectrum were extracted for each ROI and participant.

### Statistical procedures

All analyses were performed at the single-subject level before applying standard nonparametric statistical tests at the group level (Wilcoxon–Mann–Whitney unpaired tests).

### Data availability

The data that support the findings of this study are available on request from the corresponding author, B.M.. The data are not publicly available due to privacy/ethical restrictions (involvement of a clinical population).

## Acknowledgments

B.M was supported by grants ANR-16-CONV-0002 (ILCB), ANR-11-LABX-0036 (BLRI) and the Excellence Initiative of Aix-Marseille University (A*MIDEX).

## Author contributions

J.G. and B.M. designed research; A.T., P.M. and C.L.C. acquired data; J.G. analyzed data; and J.G., A.T., D.S., P.M., C.L.C., D.P. and B.M. interpreted data and wrote the paper. All authors gave comments on the paper during the process.

---

***Figure S1: Evoked activity in response to (A) 0.5 kHz and (B) 1 kHz pure tones in auditory areas. Inter-hemispheric comparison of the ITC spectra in response to pure tones, for PAC, SAC and AAC. Shaded areas indicate SEM across participants. (C-D)* Inter-hemispheric comparison of the spectral distribution of the two main ITC peaks in response *to (C) 0.5 kHz and (D) 1 kHz pure tones*, in auditory areas. Same conventions than in Figure 3.**

